# Imaging Aβ aggregation by liquid-phase transmission electron microscopy

**DOI:** 10.1101/2024.01.29.577710

**Authors:** Gabriel Ing, Silvia Acosta-Gutiérrez, Michele Vendruscolo, Giuseppe Battaglia, Lorena Ruiz-Pérez

## Abstract

The amyloid beta peptide (Aβ) readily aggregates into amyloid fibrils. This process has been the subject of intense investigations since it is associated with Alzheimer’s disease. However, it has been highly challenging to observe the microscopic steps in the aggregation reaction directly and to characterize the oligomeric assemblies formed as intermediates. To achieve this goal, we apply liquid-phase transmission electron microscopy (LTEM) in combination with all-atom molecular dynamics simulations. Our results offer an initial visualization of the dynamics of Aβ oligomers, the formation of Aβ protofibrils, and the presence of Aβ oligomers on the surface of Aβ fibrils. This work illustrates how the application of LTEM to the study of protein aggregation in solution enables the imaging of key molecular events in the aggregation process of Aβ.

## Introduction

The assembly of Aβ into amyloid fibrils and amyloid plaques is associated with Alzheimer’s disease (1, 2). While many questions remain open about the origins of the pathological processes causing Alzheimer’s disease, including the exact mechanisms of neurotoxicity (1, 2), an accurate characterization of the aggregation pathway of Aβ could provide valuable insights for the development of diagnostic and therapeutic tools for this disease (3–5).

Aβ monomers are disordered in solution (6), and also in the amyloid state, both the highly positively charged N-terminus and the hydrophobic C-terminus remain conformationally heterogeneous, forming a fuzzy coat in the amyloid fibrils (7). The monomers of both the 40-residue (Aβ40) and the 42-residue (Aβ42) forms of Aβ have a molecular mass of just over 4 kD, with an overall dimension of approximately 1 nm (8). Aβ monomers are poorly soluble and tend to self-associate to form globular structures known as oligomers (9, 10). Populations of Aβ oligomers with a wide variety of different sizes and structural features have been observed (4, 11–15). These species tend to dissociate rapidly, but in some cases, they can convert into more ordered structures and grow into amyloid fibrils (10). The generated fibrils can elongate by adding monomers at the fibril ends, a process known as elongation (3). Furthermore, amyloid fibrils can act as catalysts for further nucleation, thereby increasing the rate of oligomerization at the fibril surface, a process known as secondary nucleation (16, 17). Secondary nucleation dramatically increases the concentration of both oligomers and fibrils via a positive feedback loop. The molecular mechanisms responsible for secondary nucleation are still unknown, but recent investigations based on chemical kinetics (16–18) suggest that secondary nucleation may occur in a two-step process, with the attachment of two or more monomers to the fibril surface followed by a conformational change of the monomers to form an aggregate (18).

A wide range of methods, including cryo-electron microscopy (cryo-EM), mass spectrometry, nuclear magnetic resonance (NMR) spectroscopy, and atomic force microscopy (AFM), have been used to investigate the structures populated during the process of Aβ oligomerization, fibril formation, elongation and secondary nucleation. Through a series of recent studies, cryo-EM has proven highly effective for determining the structures of amyloid fibrils (19, 20), and also relatively homogeneous sub-populations of oligomeric species (21, 22). However, it remains challenging to use this method for investigating oligomer populations, which exhibit a high degree of conformational heterogeneity. Mass-spectrometry has been used to determine the stoichiometry of oligomers (11). NMR spectroscopy can effectively probe Aβ aggregation at high resolution, yielding real-time atomic-resolution details of the process (23, 24). AFM is an effective technique to probe structures on a surface (25), and can also be performed in a liquid environment (26). Fibrils can be examined at sub-nanometer resolutions, with a high resolution for fibril depth (27–30). We also note that computational methods, including molecular dynamics, can be helpful in understanding the aggregation process. However, these approaches are currently limited by the timescales of the aggregation processes, which is far beyond what is achievable with all-atom simulations (31). This limitation can be largely overcome by using coarse grain models, which can provide useful information on at least certain aspects of the process (10).

In this study, we use liquid-phase electron microscopy (LTEM) to image the aggregation process of Aβ. LTEM is a rapidly developing technique in which a sample is placed in a liquid cell sandwiched between two thin silicon nitride or graphene membranes. LTEM allows imaging of fully hydrated and freely evolving samples (32), and has been employed for investigating inorganic processes, including nanoparticle formation (33) and crystallization (34, 35). Although biological uses of LTEM have thus far been limited (36–43), the technique offers tremendous potential in organic and biological systems where dynamics in solution play a critical role. This potential is due to the ability to combine structural and dynamic information, which can be reported through high-resolution videos.

## Results and Discussion

### Sample preparation and characterization

Aβ40 oligomers and fibrils were prepared using previously published protocols (44, 45), and imaged by both negative-stained solid-state TEM and LTEM (**Figure 1**, and Supporting Information). Individual Aβ40 oligomers in these samples range from approximately 2 to 70 nm in diameter. This extensive size range was consistently seen across many samples at different time points along the aggregation reaction.

**Figure 1.**
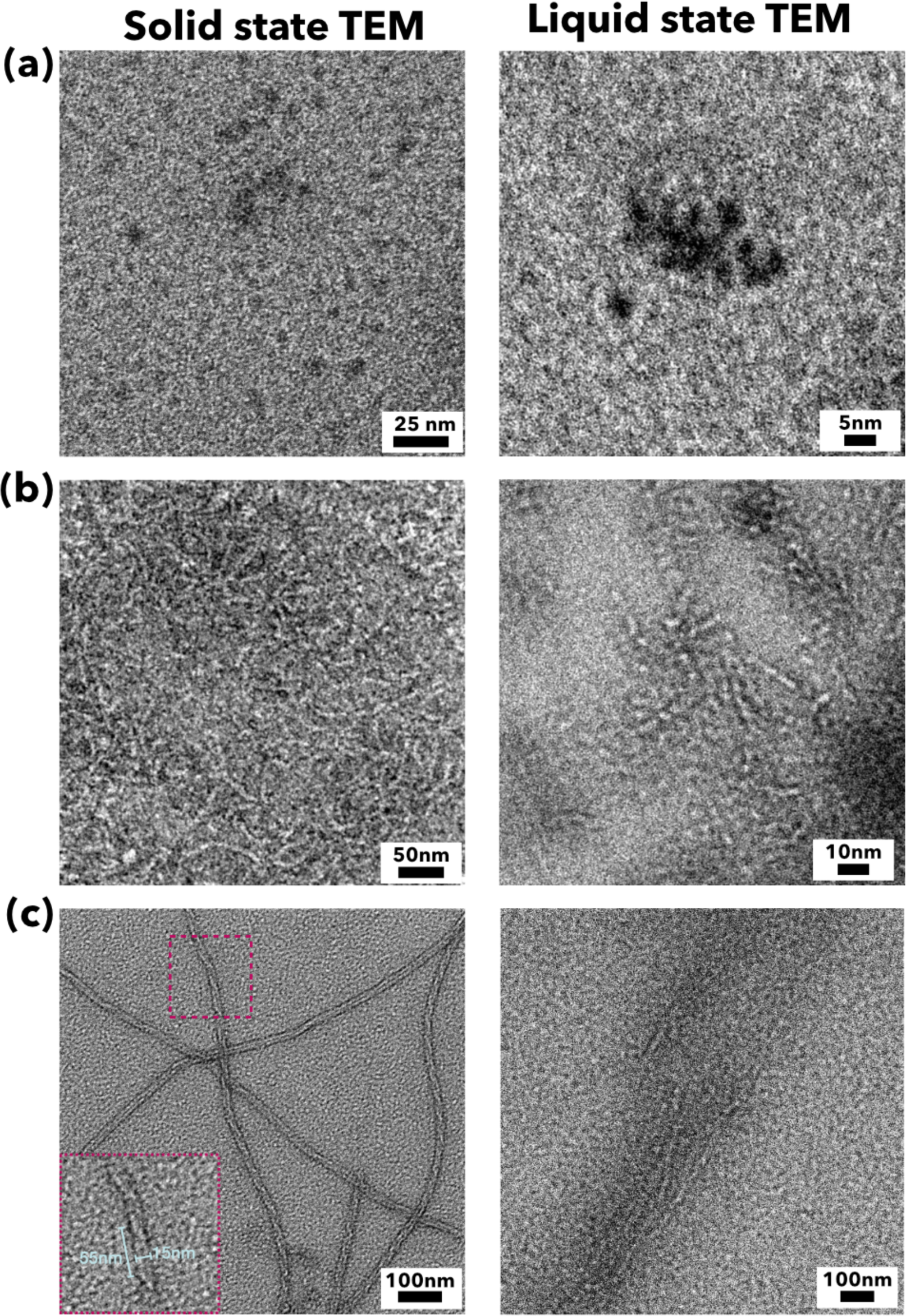
Representative micrographs from negative-stain TEM (left) and liquid-phase TEM (right). **(A-C)** These micrographs show small oligomers **(A)**, protofibrils **(B)**, and mature fibrils **(C)**.

The structures observed in fibril-forming Aβ40 samples changed significantly over time. After 1 h, only small Aβ40 oligomers were observed (**Figure 1A)**. Many small protofibrillar structures were present after 24 h of sample preparation (**Figure 1B)**. After a few days, the system seemed to reach an endpoint in structural changes within the fibril morphology, with the appearance of long mature fibrils. The fibrils varied in morphology, with thickness ranging from 7 to 15 nm, with differences in twist also observable (**Figure 1C)**. The imaged fibrils also presented high length-to-diameter ratios, as also reported by the literature (46, 47).

The comparison between TEM and LTEM results is consistent with recent findings that indicate that sensitive specimens in liquid water environments can withstand higher electron dose tolerance than when immersed in vitrified water environments, as in the case of cryo-TEM (48, 49). Such enhancement in tolerance to the electron dose likely lies in the higher diffusion rates of free radicals in liquid compared to vitrified media. Furthermore, the current investigations in liquid-phase TEM employ low-electron flux conditions for imaging protein samples to minimize damage.

### Imaging the dynamics of Aβ40 oligomers in solution

Aβ40 oligomers were imaged in a sample incubated at 4 °C for ten days. At this time point, under the conditions used here, the sample exhibited a broad distribution of oligomer sizes. To investigate the conformations of the oligomers present in the field of view, a dose fractionation video (**Video 1**) was acquired. This video was aligned and averaged, as shown in **Figure 2A**.

**Figure 2.**
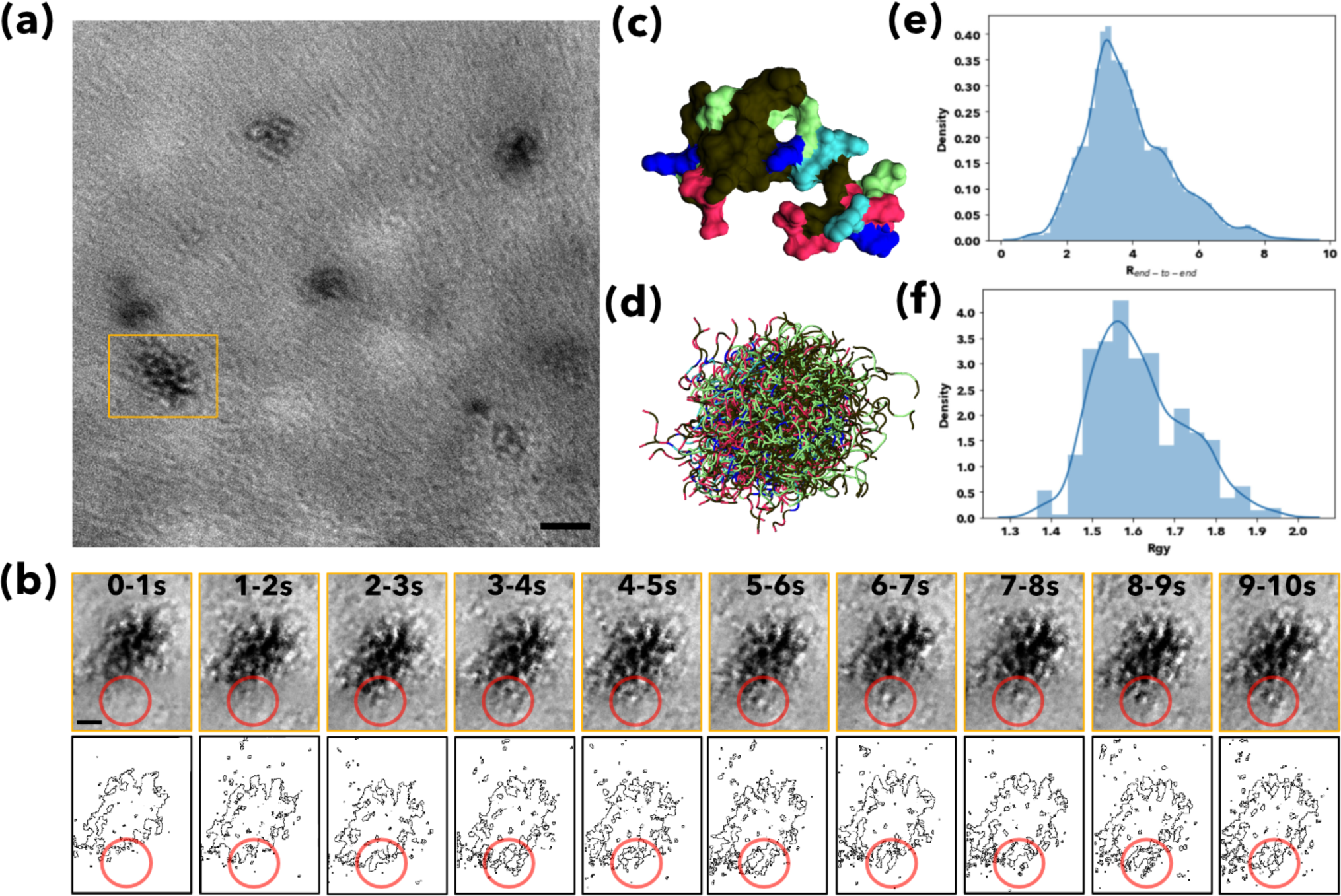
Liquid-phase TEM imaging of Aβ40 oligomers dynamics. **(A)** Full field of view of Video 1, showing large and small Aβ40 oligomers, and out of focus features in the background (100 frames, 10 s sum, motion corrected). **(B)** Time series of the dynamics of an individual large Aβ40 oligomer, which is seen to change and grow over time (red circle). Frames from video 1 are summed in groups of 10 frames, giving an effective time resolution of 1 s. Scale bars are 10 nm; images were denoised using Topaz. **(C)** Molecular model of an Aβ42 monomer coloured according to the residue properties (black-hydrophobic, green-polar-blue-basic, red-acidic, histidines-cyan. The N-terminal and C-terminal Cα atoms are shown as spheres. **(D)** Structural ensembles of 6000 conformations obtained from the molecular dynamics simulations. **(E,F)** Density distribution of an Aβ42 monomer during the molecular dynamics simulations of the end-to-end distance **(E)** and of the radius of gyration **(F)**.

The oligomers appear to be changing in shape and density in time (**Video 1** and **Figure 2A,B)**, suggesting that they have highly dynamical structures. An illustration of this process is shown in **Figure 2B**, which reports the time evolution of a large oligomer of approximately 35 nm in diameter. The structure has been rotated by 90 degrees with respect to the orientation shown in **Figure 2A**. During the 10 s video, this oligomer grows, with an additional 10 nm section attaching to the structure (**Figure 2B, red circle**). Free monomers or oligomers attaching to the large oligomer cannot be resolved clearly due to fast motion, low contrast and small size, making them difficult to image at our resolution. Another example of an oligomer changing shape and internal structure over time is shown in **Supplementary Figure 1**.

### Molecular modelling of the dynamics of Aβ42 monomers and oligomers

To facilitate the interpretation of the LTEM images described above, we performed all-atom molecular dynamics of the self-assembly process of Aβ42 peptides. Starting from Aβ42 in its monomeric state (**Figure 2C)**, we monitored first the dynamics of Aβ42 monomers. Among all kinds of Aβ isoforms (50), Aβ40 and Aβ42 are believed to be the most relevant ones in Alzheimer’s disease. Although these two forms of Aβ differ only in two amino acid residues, Aβ42 exhibits higher aggregation propensity due to the two extra hydrophobic amino acids at its C-terminal (50–52). To observe aggregation at time scales accessible to molecular dynamics simulations, we used Aβ42 due to its faster kinetics in all the simulations presented in this work.

We observed that the monomeric peptides remain mostly unfolded, thus creating an ensemble of conformations (**Figure 2E)** resembling the dark round objects imaged (**Figure 2B**, 0.1 s). The end-to-end distance of the monomer during dynamics shows a broad distribution centered at ∼3.5 nm (**Figure 2F)** with an average radius of gyration of ∼1.6 nm (**Figure 2G)**, compatible with the dark objects observed in the LTEM videos. We then analyzed the oligomerization process by performing molecular dynamics simulations with two peptides (**Supplementary Figure 2**) and four peptides (**Supplementary Figure 3**). Representative snapshots for the different assembled structures sampled during dynamics are shown in **Supplementary Figure 3**. We computed the number of hydrogen bonds formed between chains along the simulation, the radius of gyration (R_G_) and the main interactions retained during dynamics. The two-peptides simulation explores aggregated structures exhibiting one to ten hydrogen bonds (**Supplementary Figure 2A**). During the simulations, we observed different conformational rearrangements (**Supplementary Figure 2E-F**), resulting in a wide distribution for R_G_ with an average value of 2.4 nm. Adding more Aβ42 units to the simulation box follows the same trend. The R_G_ value for the four-peptide simulations (**Supplementary Figure 3**) ranged from 2 to 4 nm (**Supplementary Figure 4**).

### Imaging the process of formation of Aβ40 protofibrils

We also observed other morphologies in the vicinities of the oligomers (**Video 1** and **Figure 2**). In **Video 1,** we focused on the central oligomer structure, whose diameter was about 20 nm (**Figure 3A,B)**. We observed round species of about 5 nm joining together and forming a short, straight fibril roughly 5 nm in width and 20 nm in length (**Figure 3**). The process occurred within the first 3 s of the video. However, the fibrils appear to change shape throughout the rest of the 10 s video (**Video 2** and **Figure 3C)**. To obtain these images, we used a procedure to align and average frames using the MotionCor2 method (53), which corrects for global (i.e., whole frame) motion (**Video 1**). Short fibrils were seen in the motion correction of the entire video (see Methods), after which a rolling-average motion-corrected video was created and called **Video 2**. To make this rolling average, the video was motion-corrected and then averaged into groups of 10 frames in a sliding window fashion to complete 90 consecutive frames. This resulted in an effective acquisition time of one second. This process significantly increased the signal-to-noise ratio while also retaining some time resolution. To extract more information from **Video 2**, we also used a deep-learning-based denoising method to improve the SNR. After testing various alternatives, we chose Topaz denoising (54), based on the noise2noise framework (55), for motion-corrected videos. This same process has been observed in other videos taken on other samples (**Supplementary Figure 5)**

**Figure 3.**
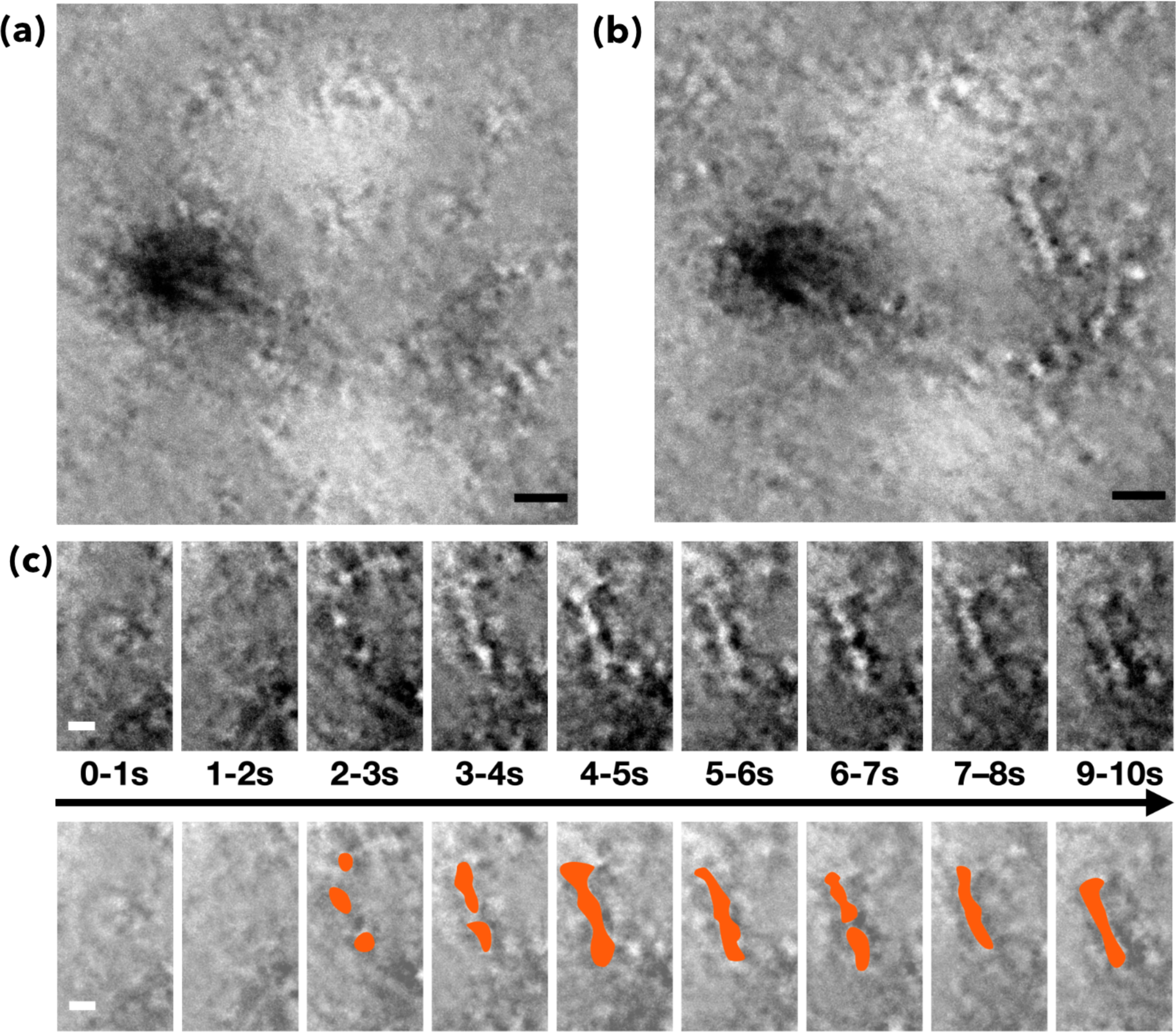
Liquid-phase TEM imaging of the formation of Aβ40 protofibrils. **(A)** Crop of 1 s motion-corrected **Video 1** frames from 0-1 s denoised with Topaz, and called **Video 2**. A large (∼20 nm) Aβ40 oligomer is observed with some blurred small structures surrounding it. The motion correction process removes objects that move rapidly between frames, resulting in images that appear blurred. **(B)** The same area as displayed in (A) but 9 s later (9-10 s), clearly displaying a short fibril that has formed and appears to be connected to some surrounding species. **(C)** Process of the protofibril formation displayed in (B) as a series of consecutive 1 s motion-corrected averages with a protofibril forming through what initially seems a pearl necklace arrangement of small monomers as shown in the third frame. Consecutive averaged frames show how the various units rearrange over time to generate the resulting short protofibril. The lower panel shows the same process as the top panel with some colouring to sketch how the process evolves over time. Scale bars are 10 nm for **(A,B)** and 5 nm for **(C)**.

### Imaging Aβ40 oliomers on the surface of the Aβ40 protofibrils

A fibril-forming Aβ40 sample was imaged in liquid after 24 h of growth at 37 °C (**Figure 4A-D)** Few fibrils could be individually identified and examined in more detail (**Figure 4B)**. One of the fibril structures appears to have a two-ply morphology, with at least two points of bifurcation where two different fibrils appear to be wrapping together (**Figure 4B)**. The captured fibrils appear to have diameters of around 3.5 nm for a single-ply fibril and 5-6 nm for a two-ply fibril (**Figure 4B,D)**.

**Figure 4.**
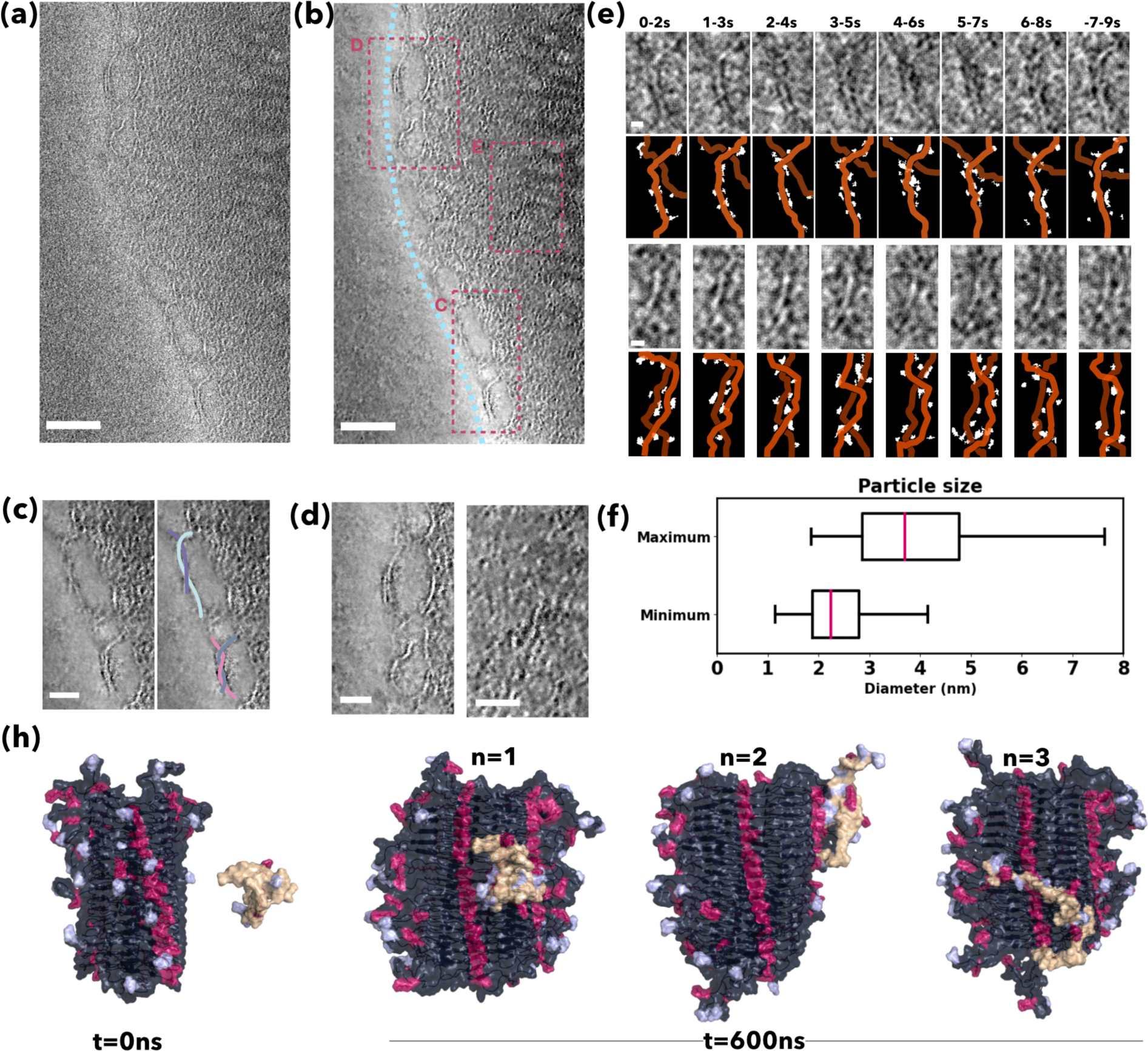
Liquid-phase TEM imaging of Aβ40 oligomers on the surface of Aβ40 protofibrils. **(A,B)** 10 s motion correction of **Video 3** before (A) after (B) denoising (**Supplementary Figure 9**), showing an uneven distribution of protofibrils within a large clotted mass on the right-hand side of the image. Areas C-E have been selected as representatives of different regions in the field of view (FOV). **(C)** Enlarged image of the interfacial area showing protofibrils twisted together forming a two-ply fibril. The fibril is traced in the right-hand image in cyan and light pink, blue and purple colours to identify the single structures forming the two-ply fibrils. **(D)** Enlarged image of the interfacial area displayed in **(B)** showing fibrils in detail. **(E)** Imaging of Aβ40 oligomers on the fibril surface. Two regions containing protofibrils, top, and bottom, are cropped from regions **C** and **D,** respectively. The frames shown are 20 s motion-corrected denoised averages. Small oligomers on the surface of fibrils are highlighted in white. These small species are shown to localise to the surface of fibrils highlighted in orange with slower dynamics and higher concentrations than the surrounding areas. **(F)** Box plot of the height and width of the Aβ40 oligomers on the surface of the Aβ40 protofibrils. Scale bars in the first frames of A and B are 5 nm. **(H)** Aβ42 monomers (orange) attached to the Aβ42fibril (dark grey) in different positions, including the disordered N-terminus and the ordered surface. Charged residues are highlighted in light blue (negative) and red (positive) after 500 ns of MD simulations. The scale bars are 100 nm for **(A, B)**, 20 nm for **(C -D)** and 5 nm for **(E)**.

The specimen solution was drop-cast directly onto the bottom chip of the liquid cell, and the cell was sealed. In this experiment, dark solid masses partially blocked the window, yet many droplets of the liquid sample could be seen. Imaging was performed inside several droplets to search for fibrils or other regions of interest. A large mass consisting of tangled fibrils was seen within a large droplet and a dose fractionation video was acquired in low-dose conditions (1.83 e^-^A^-2^s^-1^, 18.3 e^-^ A^-2^ total). This magnitude of electron dose should limit beam damage to the specimen, with only limited high-resolution damage being seen (49, 56). To increase the contrast, i.e., the SNR of the imaged fibrils in the field of view, the full hundred frames of the 10s long (10 fps) video, referred to as **Video 3**, were aligned and averaged using motioncor2 as explained above.

The denoised, sliding-window motion-corrected resulting videos (**Video 3**) could then be analyzed with a specific focus on the individual fibrils located at the edges of the mass of fibrils. In this fashion, the fibril structures displayed in **Figure 4B,C** at the bottom right-hand side area were analyzed to investigate the evolution of the structures over time and shown in **Figure 4E**. These are shown as raw and segmented images with false coloring of the fibrils to highlight the presumed monomers and fibrils. Single dark assemblies of small size appear on the fibril surface and seem to grow and change over time (**Figure 4**). Based on the size distributions obtained by molecular dynamics of monomers (**Figure 2F)** and small oligomers (**Supplementary Figures 6,7,2-4**), these small assemblies appear to be monomers or small oligomers of up to 3 or 4 monomers adhering to the fibril surface. This binding may offer a glimpse of secondary nucleation processes, but it would require measurement times beyond the current 10 s to follow this process further. These small species grow and shrink over time, suggesting that the particles are in equilibrium between being adhered to the fibril surface, providing high contrast, and rapidly moving in solution, for which low or no signal is seen. However, this is evidence that small Aβ40 species stick to the surface of fibrils and form larger oligomer species, as assumed by kinetic models (18). This observation is also supported by results in negative stained TEM, in which fibrils are commonly seen coated by oligomers (**Supplementary Figure 8**).

## Conclusions

We have reported the use of LTEM to study the aggregation process of Aβ in real-time and liquid environment, thus providing information about the structures and dynamics of the Aβ species populated during the process. The results that we have described provide a visualization of 3 aspects of Aβ aggregation: (1) the dynamics of Aβ oligomers, (2) the formation of Aβ protofibrils, and (3) the attachment of Aβ oligomers to the surface of Aβ fibrils. We anticipate that the use of faster image acquisition rates, and improved motion correction and denoising procedures will enable future studies to achieve a higher accuracy in the imaging of the dynamics of self-assembling Aβ species in solution.

## Materials and Methods

### Aβ40 peptide preparation

Aβ40 peptide (Bachem 4014442.1) was prepared into oligomer and fibril samples using previously described methods (44, 45, 57). The samples were characterized first using conventional solid-state TEM to confirm that the oligomers and fibril structures had formed as expected. The oligomer and protofibril samples shown in **Figure 1** were one day old, while the mature fibrils had aged for over a month. At that point, there was a stable population of fibrils with no obvious negative effect.

### Transmission electron microscopy (TEM)

All samples were imaged at an Aβ peptide concentration of 0.5 mg/ml. The solid-state TEM images displayed in **Figure 1** and **Supplementary Figure 8** were prepared by first drop casting 3 µl of sample onto 300 mesh-copper grids with a carbon film, waiting for 1 min, then blotting off excess sample followed by placing in a droplet of UranyLess stain, then again blotting off the excess before drying with a vacuum pump.

For the imaging performed via LTEM, DENS solutions liquid-phase holders were used, with the data presented all being prepared from the STREAM system, except for the LTEM image **Figure 1A**, which was imaged with the DENS Ocean system. The STREAM system is superior, as thinner liquid cells can be created with negative pressure from a vacuum pump. A drop-casting method was used for both these systems, with 1.0-2.5 µl pipetted directly onto the bottom chip. These chips had perpendicular window configuration and 0 nm spacers and were untreated before use. The total electron doses used for the LTEM images were 35.3 e^-^A^-2^ (**Figure 1A)**, 105 e^-^A^-2^ (**Figure 1B)**, 0.401e^-^A^-2^ (**Figure 1C)**, 170 e^-^A^-2^ (**Figures 2, 3** and **Videos 1, 2**) and 18.3 e^-^A^-2^ (**Figure 3** and **Video 3**).

A JEOL JEM2200 FS microscope equipped with a field emission gun and low-dose Gatan K2-IS camera was used for the TEM imaging. Electrons were accelerated to 200 keV. All images and videos were collected in counted mode using Digital Micrograph v.3.4, and the videos were recorded in dose-fractionation mode. Images were obtained at a rate of 10 frames per second. The images were also recorded as dose fractionations with automatic motion correction, generating a final image with reduced motion blur. The images were saved as 32-bit images in Gatan dm3 or dm4 format.

Imaging Aβ in solution is challenging, as the specimen is unstained and has low contrast due to its low atomic number. In addition, liquid cells are typically thicker than the conventional grids used for state solid-state TEM. In this regard, the electron transparent silicon nitride (SiN) windows in the liquid cell tend to bulge when inserted into the microscope due to the differential pressure between the microscope column and the cell (58). This bulging effect generates an increased thickness in the liquid media where the specimen is dispersed, which causes an increase in noise due to inelastic and multiple electron scattering events. Such scattering events obscure the signal-to-noise ratio (SNR) in the acquired images of low-contrast Aβ specimens.

Since the particles to be observed are also mobile, increasing the signal by using longer exposure times is difficult. However, alignment and averaging of video frames are often highly useful, as detailed below. The challenges presented with low contrast limitations also make the imaging process arduous when visualizing features of interest during live imaging. Biological systems imaged by LTEM, and in the present case Aβ, require careful image post-processing protocols to denoise, deblur and unveil features that are not readily discernible to the naked eye (59–61).

### Image processing: Motion correction

LTEM movies were processed using the Python package SimpliPyTEM (62). Videos were aligned using motion correction with Motioncor2 (53) to form an aligned sum of the frames from the videos with reduced motion blur. The motion correction processing was performed as a single, full-frame alignment, so only the global motion, either from stage drift or scale large-scale liquid movement in the imaging area, is corrected. Motion correction was performed in groups of 10 or 20 frames in a sliding window across the 100-frame videos of 10 sec to retain dynamic information while increasing the signal-to-noise ratio.

### Image processing: Denoising

To denoise corrected motion movies, we used the neural network-based denoiser Topaz (54). While attempts were made to train this denoiser on liquid-TEM videos, the best performance was achieved using the pre-trained default uNet model that was shipped with the software. While these models were trained on cryo-TEM images rather than LTEM images, the training datasets available to the developers were much larger than we used hence the better performance. **Supplementary Figure 9** shows the image process workflow.

### Molecular dynamics simulations

We simulated a fibril of Aβ42 in the presence of one or three Aβ42 monomers. The cryo-EM structure 7Q4B was used as the starting point to build the fibril system. This structure was extended to 32 total chains (17 pairs), and the N-terminal residues, which were not present in the original structure due to flexibility, were built onto the structure. Following the building of the fibril, it was solvated, 100 mM sodium and chloride ions were added, and the system was parameterised using the TIP4P (63) water model and Amber99-Disp force field, which has been designed to deal well with both folded and disordered structures (64).

The aggregation of Aβ42 was simulated using the same force field starting from 32 structures of the linear peptide. All simulations were performed with Gromacs v2020.6 with a 2 fs integration step. Each system was energy minimised using the steepest decent algorithm, heated with 5000 steps of NPT at 300K and equilibrated with 50000 steps of NVT. A 100 ns run of the fibril system preceded adding either one or three monomers into the system, followed by the same minimisation, NPT and NVT steps for the individual simulation. The final box dimensions and numbers of atoms were (15.79, 14.88, 12.89), 392,132 for one monomer and (17.60, 16.58, 14.38), 545,179 for three monomers. The production run for all systems was 1 µs.

## Supplementary Information

### Molecular modelling of Aβ42 monomers on the surface of Aβ42 fibrils

To facilitate the interpretation of the LTEM images, we performed all-atom molecular dynamics at an increasing number of peptides in the simulation box. Starting from the single peptide or monomer Aβ42 (**Figure 2C)**, we performed simulations using the AmberDisp force field with TIP4D water to ensure a disordered behaviour of Aβ42 in solution (64, 65). Structural ensembles of 6000 conformations were obtained (**Figure 2D**). We monitored the structural evolution during dynamics. To gain insight into the attachment process of monomers on the surface of the imaged fibrils, we performed 1 μs all-atom molecular dynamics simulations on a fibril model surrounded by either one or three Aβ42 monomers. These simulations show how Aβ42 monomers interact with a fibril surface, as this is likely a key step in secondary nucleation processes. An all-atom model of a short Aβ42 fibril was built to perform this simulation based on the cryo-EM structure (7Q4B). The N-terminal regions (residues 1-8) of the chains of these fibrils are not built in the cryo-EM structure due to inherent flexibility causing heterogeneity in the cryo-EM populations; these were manually built into the structure with PyMOL. Although these two kinds of Aβ differ only in two amino acid residues, Aβ42 exhibits higher aggregation potential due to the two extra hydrophobic amino acids in its C-terminal (54–56). To observe aggregation at molecular dynamics reachable time scales, we decided to test Aβ42 due to its faster kinetics in all the simulations presented in this work.

In all simulations performed (three repeats with a single monomer, three replications with three monomers), the monomers are found to adhere to the surface of the fibril (**Supplementary Figure 7**). This adherence is seen across all simulation runs (with either one or three monomers) for an average of >80% of the frames (**Supplementary Figure 7**B). The majority of interactions seen are found towards the N-terminus of the monomer (**Supplementary Figure 7C**). This behaviour could allow oligomerisation as the hydrophobic C-terminus is free to diffuse in a small radius, which could then interact, through hydrophobic interactions, with free monomers. This observation is consistent with the possibility that the species seen moving on the surface of fibrils in **Figure 4** are Aβ42 monomers or oligomers.

**Supplementary Figure 1.**
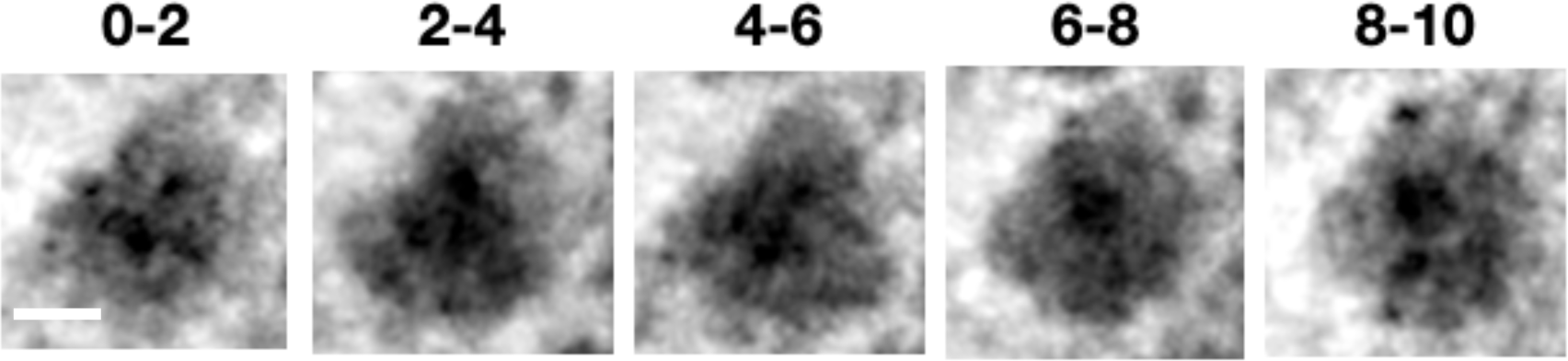
Liquid-phase TEM imaging of Aβ40 oligomers dynamics. (A) Cropped area of view of **Video 4**, showing a 25nm Aβ40 oligomer (100 frames, 20 s sum, motion corrected). Time series of the dynamics of the oligomer, which is seen to change shape and rearrange its internal units over time. Frames are summed in groups of 20 frames, giving an effective time resolution of 2 s. Scale bar is 10 nm; images were denoised using Topaz.

**Supplementary Figure 2.**
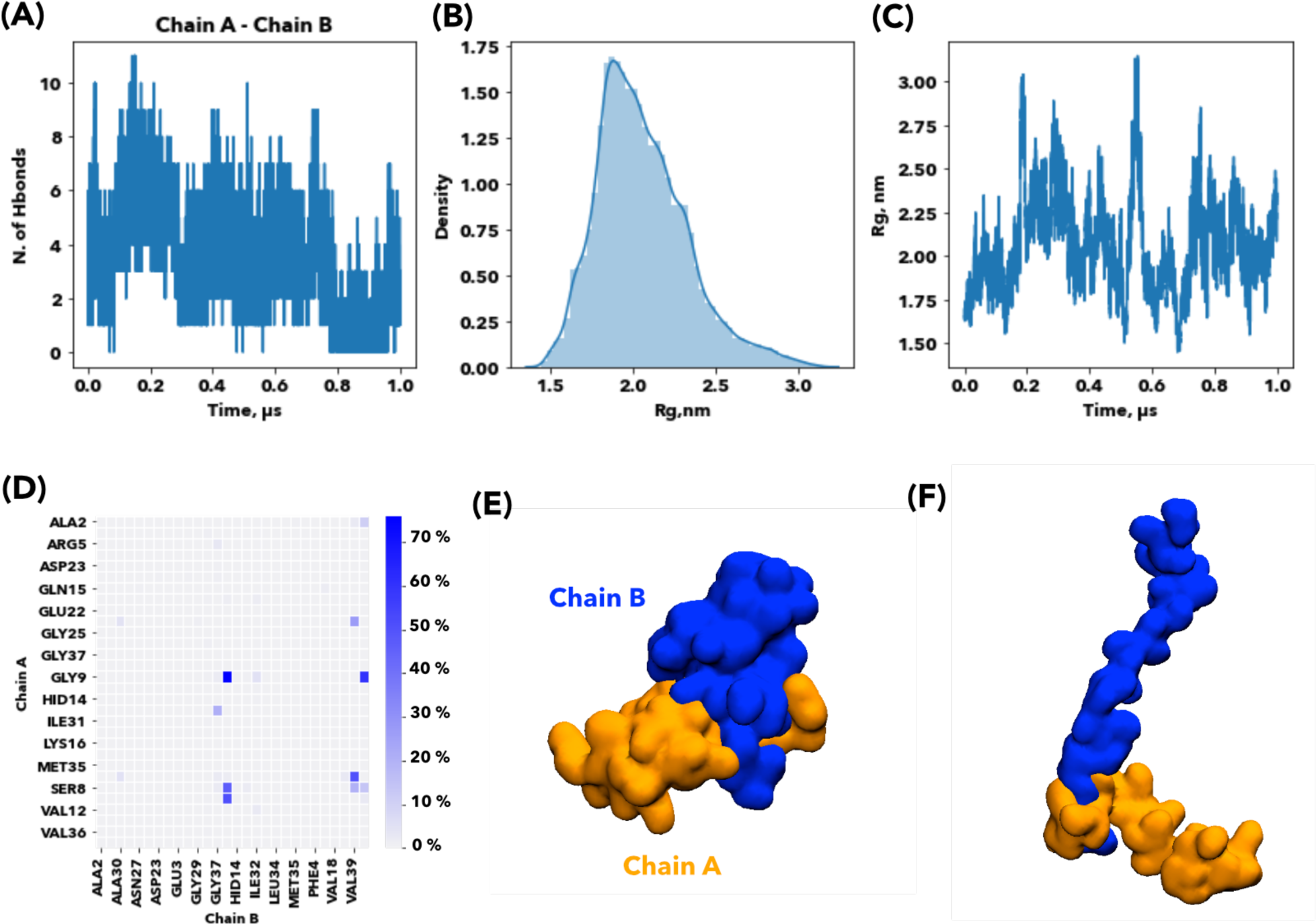
Interactions between two Aβ42 monomers in molecular dynamics simulations. **(A)** Number of hydrogen bonds formed during dynamics between two Aβ42 peptides (A and B). **(B)** Radius of gyration distribution of a total of 6600 dimer conformations. **(C)** Time evolution of the radius of gyration. **(D)** Contacts between the Aβ42 peptides during simulation are coloured according to persistence during the simulated time. **(E)** Snapshot representative of a compact dimer conformation. **(F)** Snapshot representative of a more extended dimer conformation. Each chain is represented as its van der Waals surface and coloured according to the peptide id A (orange), B (blue).

**Supplementary Figure 3.**
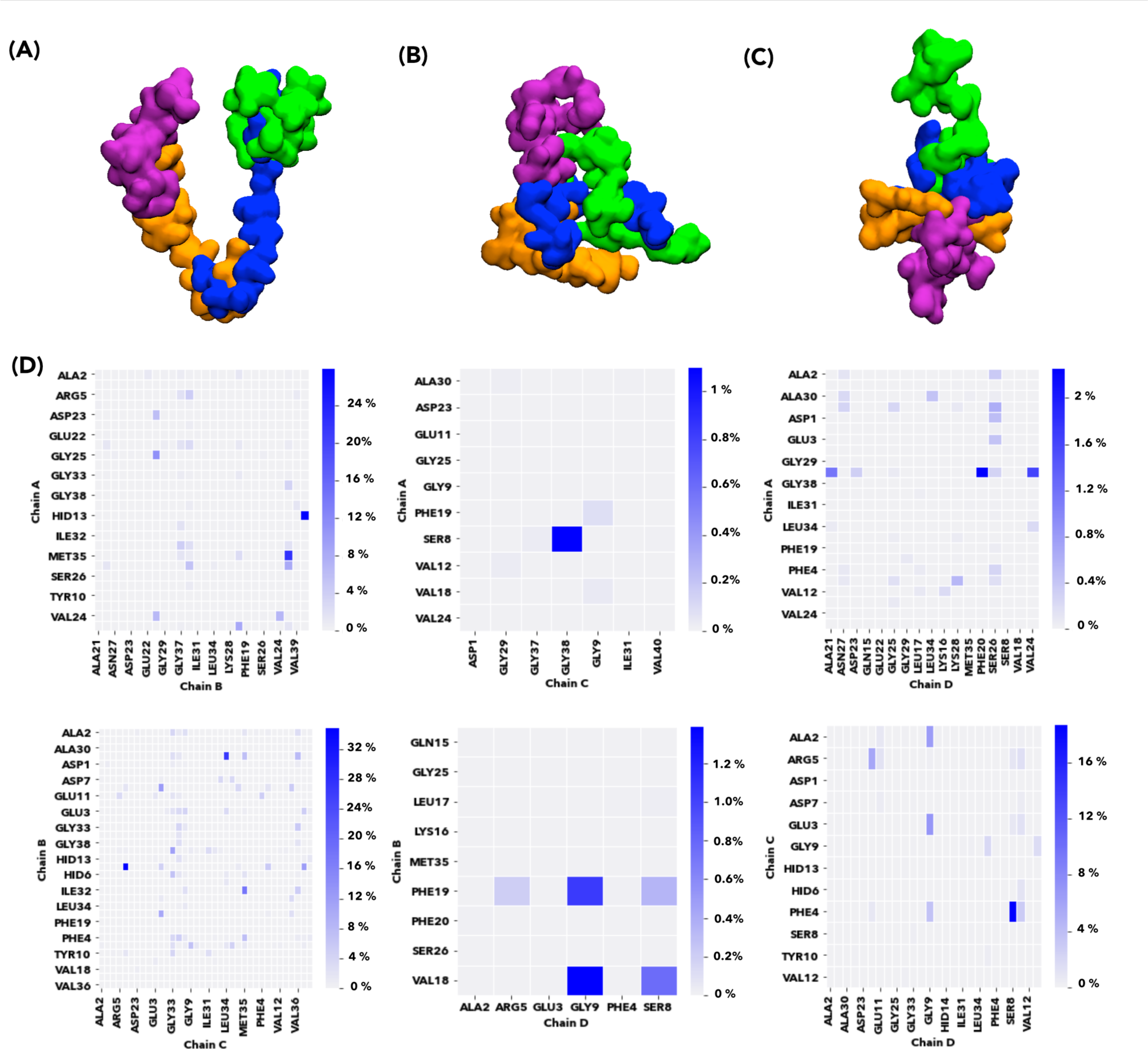
**(A-C)** Snapshot representatives for the different assembled structures sampled during dynamics of Aβ42 peptides. Aβ42 peptides are depicted as van der Waals surfaces coloured according to chain id from A to D (orange, blue, green, purple). **(D)** Contacts between Aβ42 peptides are coloured according to the percentage of the simulation in which they remain formed.

**Supplementary Figure 4.**
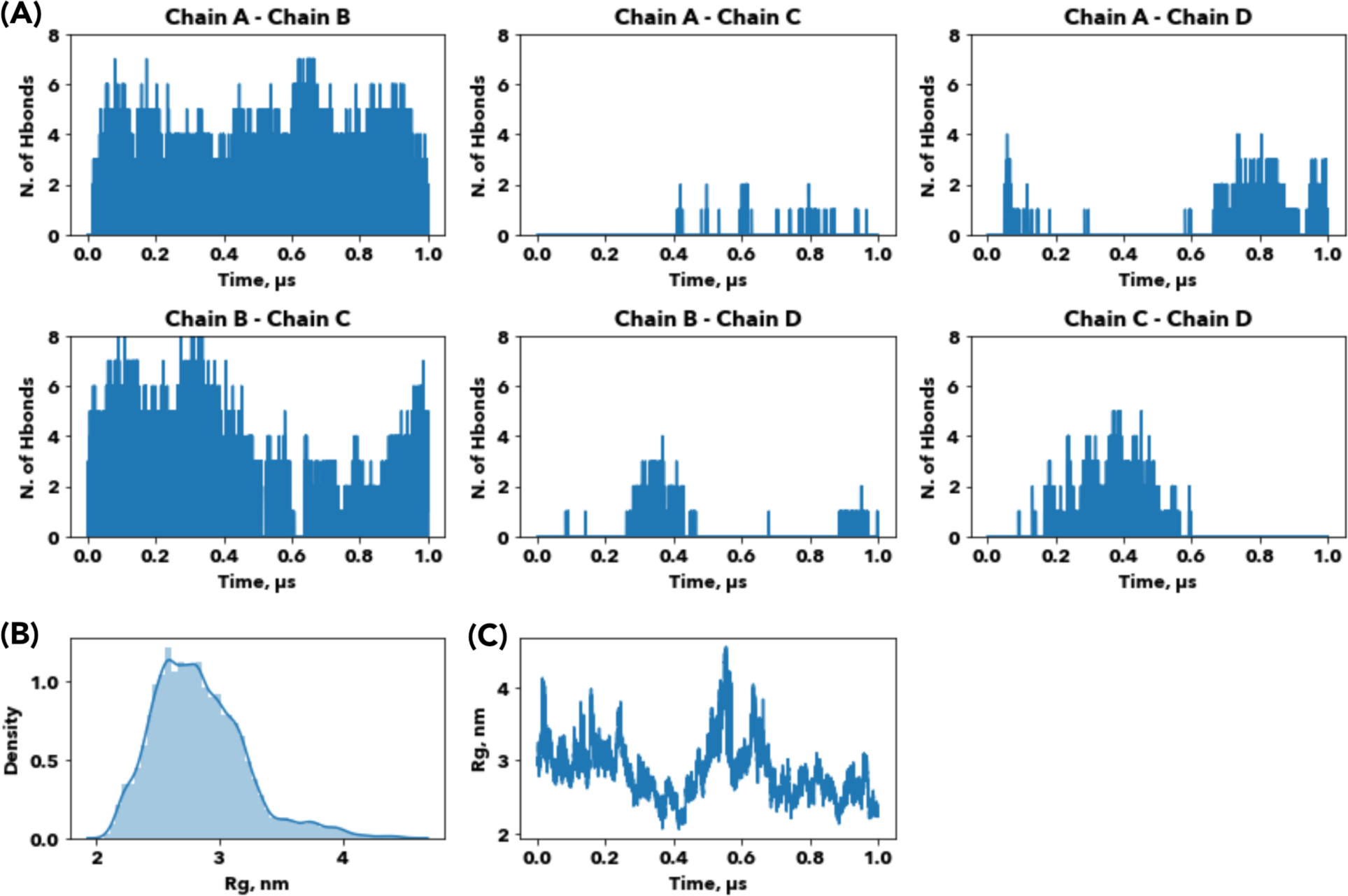
Interactions between four Aβ42 monomers in molecular dynamics simulations. **(A)** Number of hydrogen bonds formed during dynamics between the four Aβ peptides versus simulation time. **(B)** The radius of gyration distribution of a total of 50000 conformations. **(C)** The radius of gyration evolution over simulation time.

**Supplementary Figure 5.**
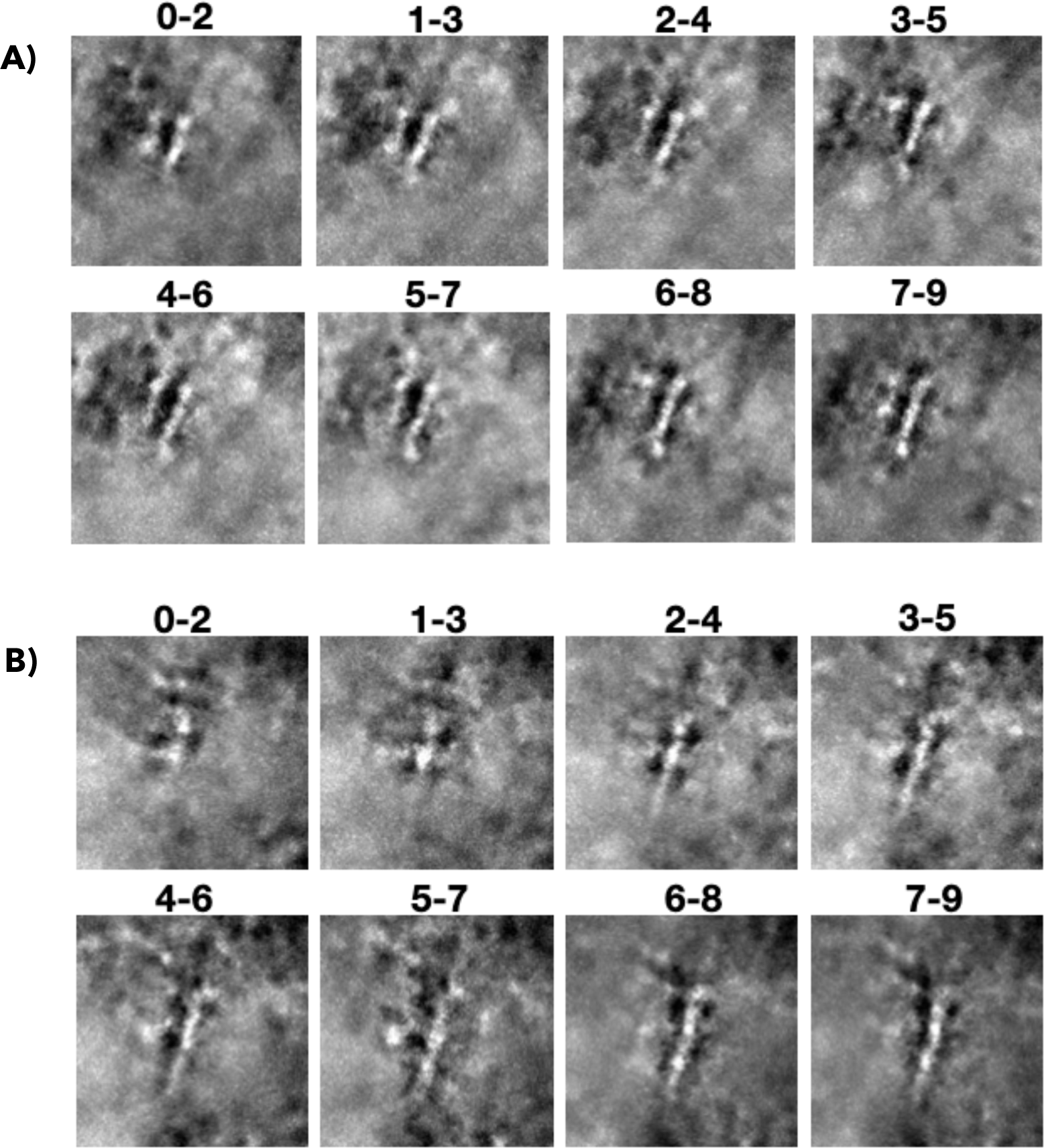
Liquid-phase TEM imaging of the formation of Aβ protofibrils. Two different areas **(A-B)** cropped of 2 s motion-corrected **Video 5** denoised with Topaz. **(A)** and **(B)** show some units assembling at the start of the video (0-2s) and 9 seconds later (7-9s), the units appear to rearrange over time in consecutive frames to generate the resulting short protofibrils. Scale bars are 5 nm for **(A,B)**

**Supplementary Figure 6.**
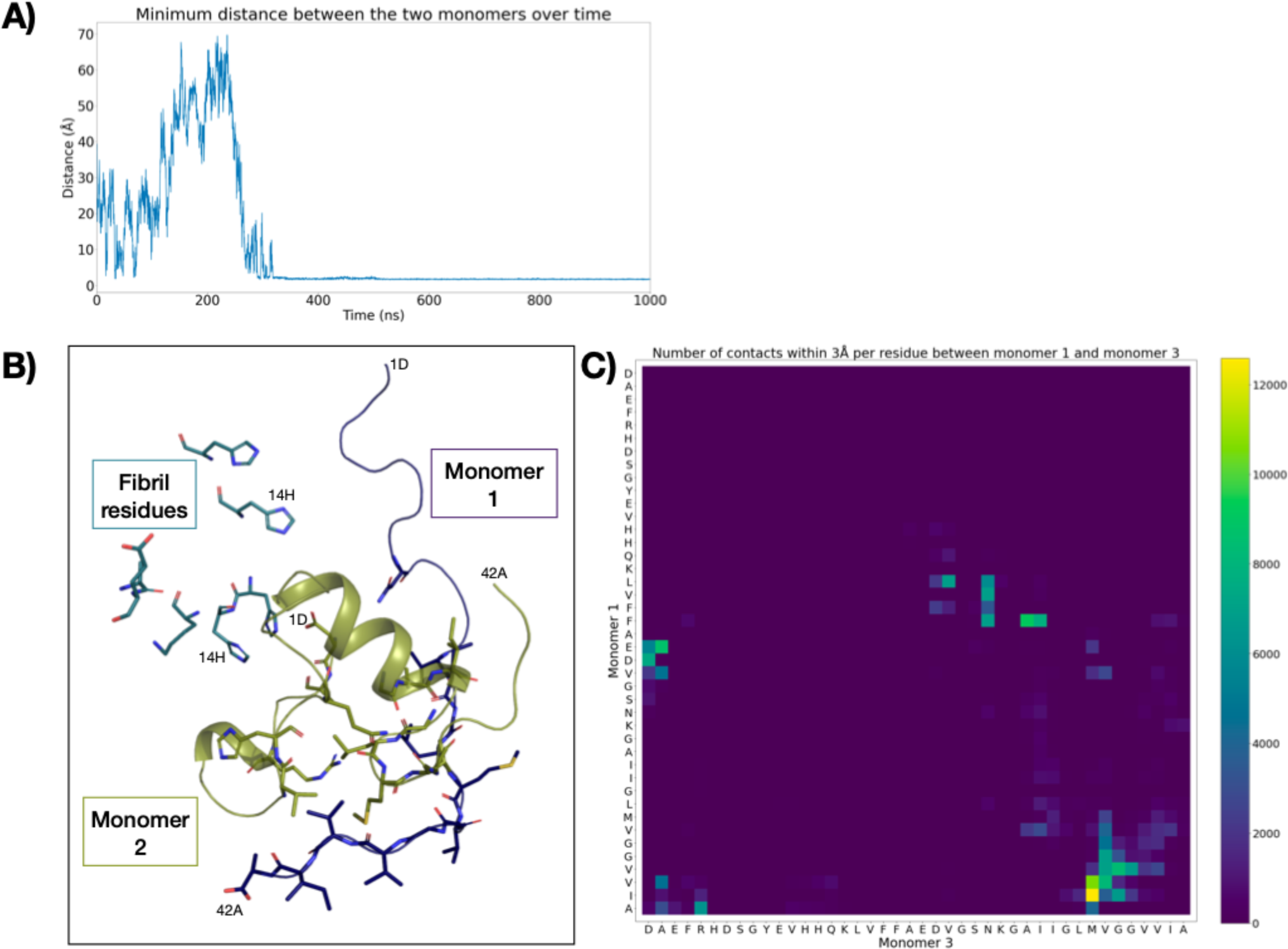
Interactions between two Aβ42 monomers on the surface of Aβ42 fibril in molecular dynamics simulations. **(A)** distance between the two Aβ monomers over time. This graph shows that once the monomers make contact in the correct way, the contact is very stable as the monomers do not separate for over 700 ns. **(B)** Snapshot of interactions at 500 ns. Monomer 1 is seen to wrap around monomer 2, which is connected to the fibril through interactions with the histidine residues at position 13 and 14 of the fibril chain. The interactions between the monomers are largely hydrophobic in nature, with the hydrophilic residues or atoms facing away from the opposite monomer. **(C)** Heat map of interactions between two monomers, showing the interaction is mainly mediated by C-terminus hydrophobic residues.

**Supplementary Figure 7.**
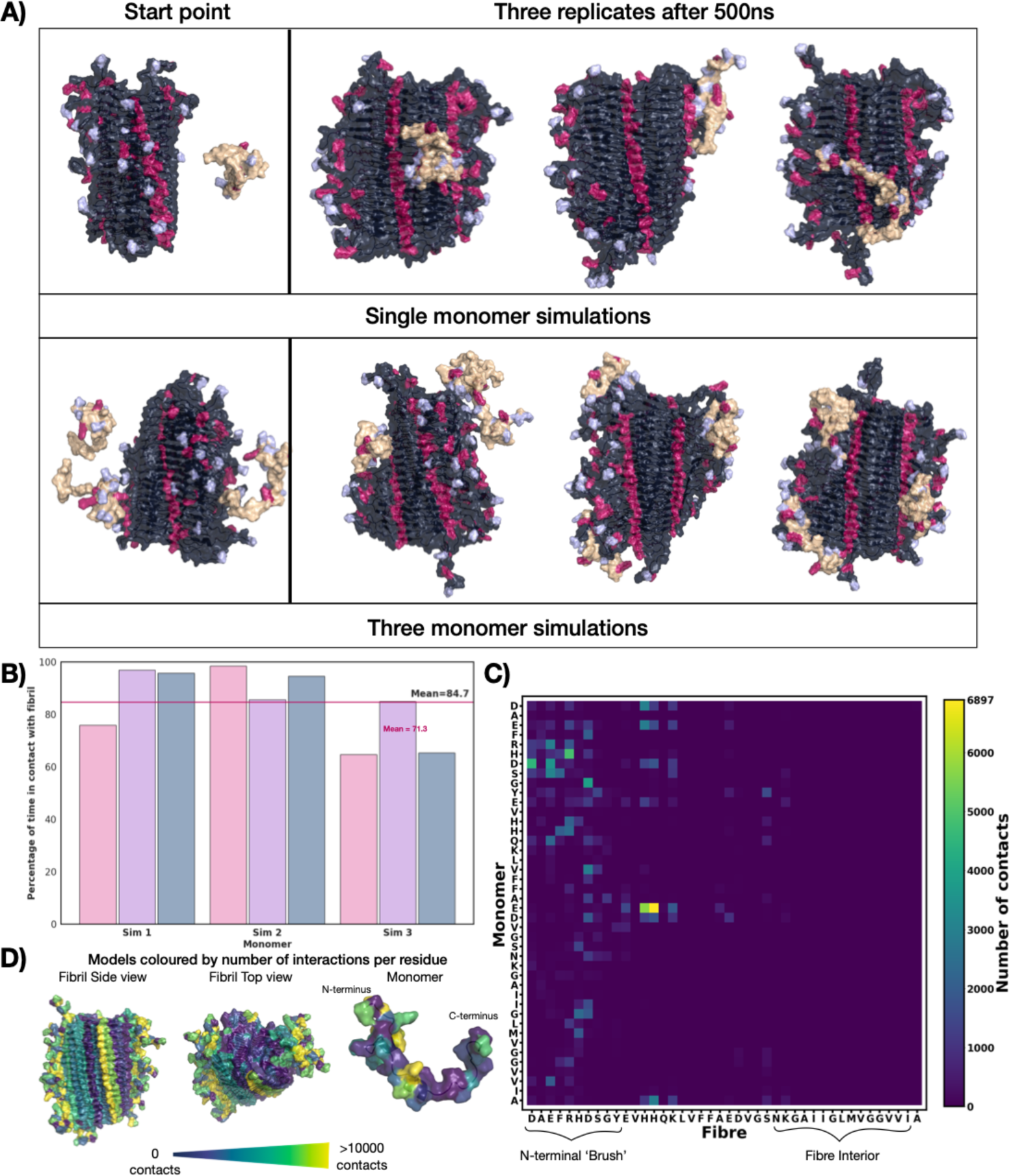
Analysis of the interactions between Aβ42 monomers and fibrils in molecular dynamics simulations. **(A)** Positions of the Aβ monomers (orange) on the fibril (dark grey) as seen in 3 repeats of MD simulations after 500 ns. The charged residues are highlighted in light blue (negative) and red (positive). Aβ monomers are seen attaching in various positions including the disordered N-terminus and the ordered surface of the fibril. **(B)** Percentage of time each monomer is in contact (< 2 Å) with fibrils in MD simulations with 3 monomers. This graph shows that for all four simulations performed (total 2.2 µs) the monomers are all in contact with the fibrils for the majority of the time sampled, with only two exceptions, one monomer was in connection for only 40% of the time, while the other never reached the fibril in the time sampled. **(C)** Heat map showing which residues on both the monomer (y axis) and fibril (x-axis) are in in contact. This heat map shows the contacts are mediated largely by unspecific interactions within the disordered N-terminus bush of the fibril, with the exception of a clear major interaction point on the ordered surface of the fibril, while on the monomer the majority of interactions are at the charged N-terminus, but contacts are seen across the rest of the peptide sequence. **(D)** Locations of the residues commonly in contact coloured on the fibrils and monomer models.

**Supplementary Figure 8.**
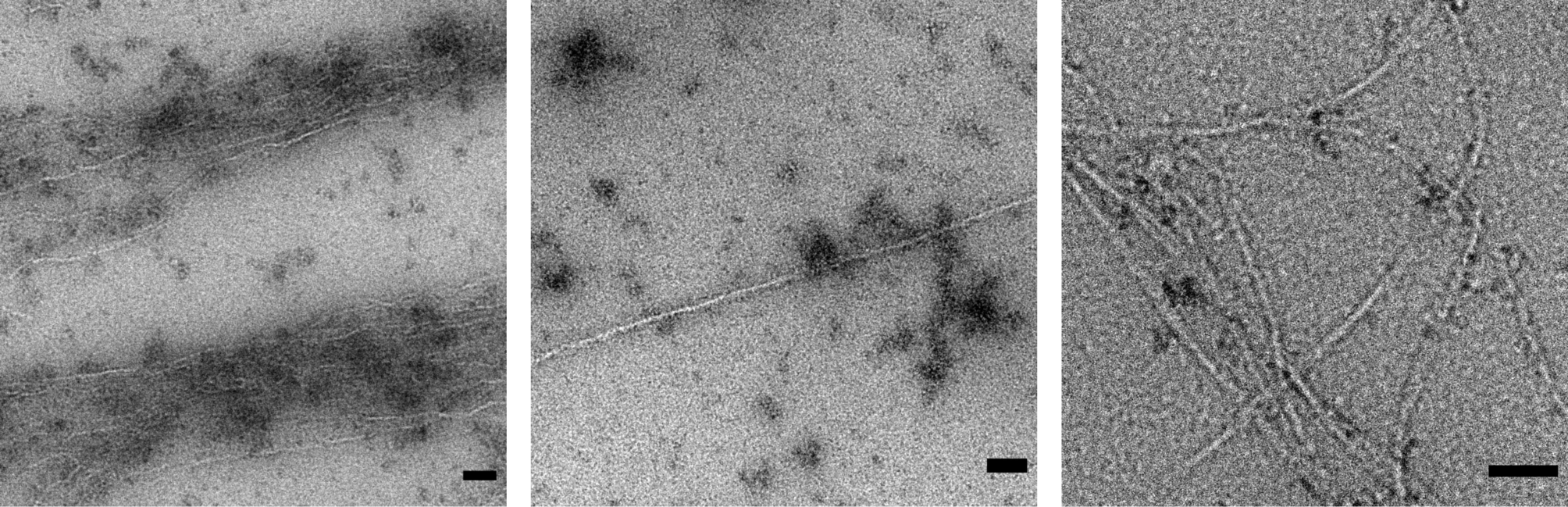
Micrographs showing Aβ40 oligomers localised to the fibril surface via dry TE M. Aβ40 monomers and oligomers are consistently localised to the surface of fibrils in dry TEM. Scale bars are 50 nm.

**Supplementary Figure 9.**
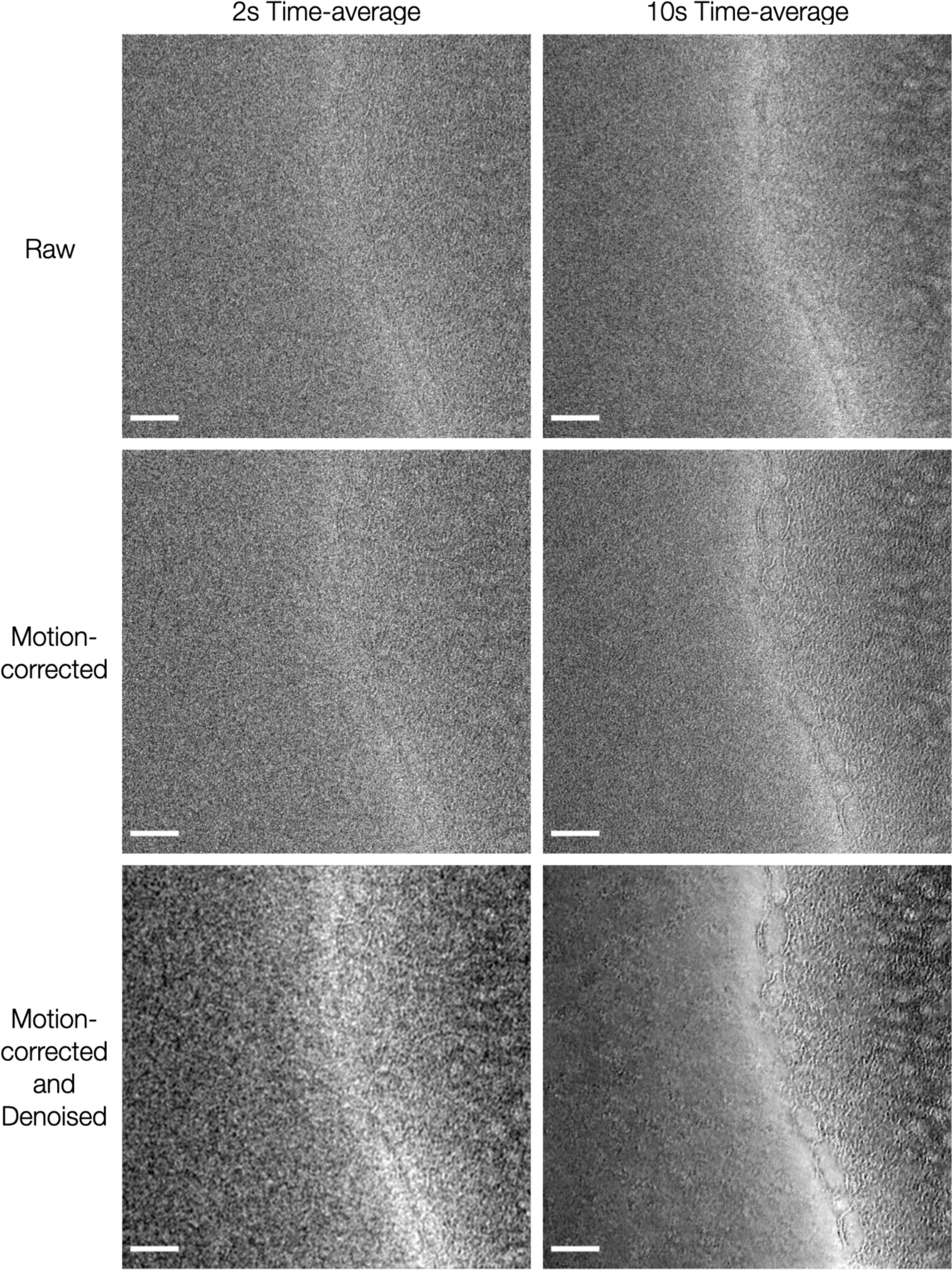
Micrographs showing the image processing workflow. Multiple frames of the 100 frames of 10 s long **Video 3** and shown in Figure 4 were summed together to decrease noise and increase signal in the images. The values chosen were full 10 s time averages for a global and sharp view of the fibrils present, and 2 s averages to retain time resolution. The raw time-averaged frames show very little contrast, however, when a motion-corrected procedure is used, the fibrils are sharper. Topaz denoising is also applied with the pre-trained uNet model, this step significantly reduces the noise, crucially making the fibrils visible in the 2 s time-averages. Details can be seen in the background of the 10 s denoised, motion-corrected time-average. Scale bars are 50 nm.

